# Biomechanical Modeling of Cesarean Section Scars and Scar Defects

**DOI:** 10.1101/2023.11.03.565565

**Authors:** Adrienne K. Scott, Erin M. Louwagie, Kristin M. Myers, Michelle L. Oyen

## Abstract

Uterine rupture is an intrinsically biomechanical process associated with high maternal and fetal mortality. A previous Cesarean section (C-section) is the main risk factor for uterine rupture in a subsequent pregnancy due to tissue failure at the scar region. Finite element modeling of the uterus and scar tissue presents a promising method to further understand and predict uterine ruptures. Using patient dimensions of an at-term uterus, a C-section scar was modeled with an applied intrauterine pressure to study how scars affect uterine stress. The scar positioning and uterine thickness were varied, and a defect was incorporated into the scar region. The modeled stress distributions confirmed clinical observations as the increased regions of stress due to scar positioning, thinning of the uterine walls, and the presence of a defect are consistent with clinical observations of features that increase the risk of uterine rupture.

## 1 Introduction

The current global Cesarean section (C-section) rate is about 21% and is increasing worldwide, especially in developing countries [1, 2]. C-sections are a major abdominal surgery requiring months of recovery time for patients and adding substantial costs to the healthcare system [2, 3]. Although C-sections are essential for decreasing maternal, neonatal, and infant mortality rates, this major surgical procedure can sometimes be performed unnecessarily [4]. In fact, the World Health Organization reports that C-section delivery rates above 10 – 15% are not justified to reduce mortality [5, 6]. Therefore, there is a need to limit the overuse of C-sections and the associated short-term and long-term complications resulting from this surgery [5].

One inevitable result of a C-section is uterine scarring from the C-section incision. Additionally, inadequate healing of C-section scars can lead to defects in the scar tissue, also known as an isthmocele or niche [6]. Both the scar and scar defects can lead to a loss of tissue mechanical integrity in a subsequent pregnancy resulting in an increased risk of uterine tissue failure known as uterine rupture or uterine dehiscence: uterine rupture is a complete division of the three tissue layers in the uterus (perimetrium, myometrium, and endometrium), while uterine dehiscence is only a partial division of the uterine layers [7]. In a subsequent pregnancy after a C-section the rate of uterine rupture increases to approximately 0.5% [8]. In comparison, the rate of uterine rupture is only about 0.02% in patients with no prior C-section [9]. Although a rate of 0.5% is low, uterine rupture is a life-threatening emergency as neonatal mortality ranges from 74 to 94% and maternal mortality ranges from 1 to 13 % due to uterine rupture [10].

To prevent uterine rupture, it is important to consider all of the associated risk factors in addition to the presence of a scar and scar defects. Clinical data has shown that the positioning of the scar alone influences the risk of uterine rupture. Historically, the first classical C-section incisions were vertical along the sagittal plane. Now, it is well known that vertical scars in the uterus further increase the risk of uterine rupture [11]. For this reason, vertical incisions are avoided unless needed for emergencies, and low transverse incisions are used instead. Recent literature has also shown how uterine structure, such as thinning of the lower uterine segment measured by 2D ultrasound, correlates to uterine rupture risk [12, 13]. Additionally, connective tissue disorders that are likely to affect the mechanical properties of the tissue are also associated with an increased risk of uterine rupture [14]. Taken as a whole, it is evident that uterine rupture is a mechanical process that could be evaluated using engineering tools.

Although some risk factors have been identified, there is currently no reliable model to predict uterine rupture. Furthermore, to our knowledge, no one has previously examined the risk of uterine rupture with C-section scars and scar defects with a biomechanical approach. Finite element (FE) modeling of the uterus and the scar tissue may present a promising method to further understand and predict uterine ruptures. FE models allow for the study of how various parameters (patient geometry, mechanical properties) influence uterine mechanics. Therefore, our objective is to generate a mechanical model of the uterus to examine how the scar positioning (sagittal versus transverse), material properties of the scar, and the presence of a defect influence uterine tissue mechanics.

## 2 Methods

### Parametric Geometries

A baseline representative model of a uterus and cervix was generated by averaging patient uterine dimensions acquired from MRI images of five at-term pregnant patients and constructing a geometry in Solidworks 2022 (Dassault Systémes, France) as previously described [15–17]. A scar (Fig. 1A) and surrounding abdomen tissue were also included in the Solidworks model [18]. The scar was split into two parts to allow variance of the material properties at the midline of both the transverse and sagittal scar (Fig. 1B). The standard deviation of four uterine thickness measurements (fundal, anterior, left/right, and lower uterine segment thickness, or UT1, UT2, UT3, UT4 respectively) of the five at-term pregnant patients were calculated and added or subtracted from the average baseline model (Fig. 1C). Geometries were then built in Solidworks to vary the uterine thickness by increasing the thickness values by a standard deviation (baseline + SD) and decreasing the thickness values by a standard deviation (baseline – SD) (Fig. 1C). Next, a defect was incorporated in to the scar tissue (Fig. 1D). Dimensions of the defect size (i.e., width, depth, length) were based on average measurements found in the literature [20, 19].

**Fig. 1.**
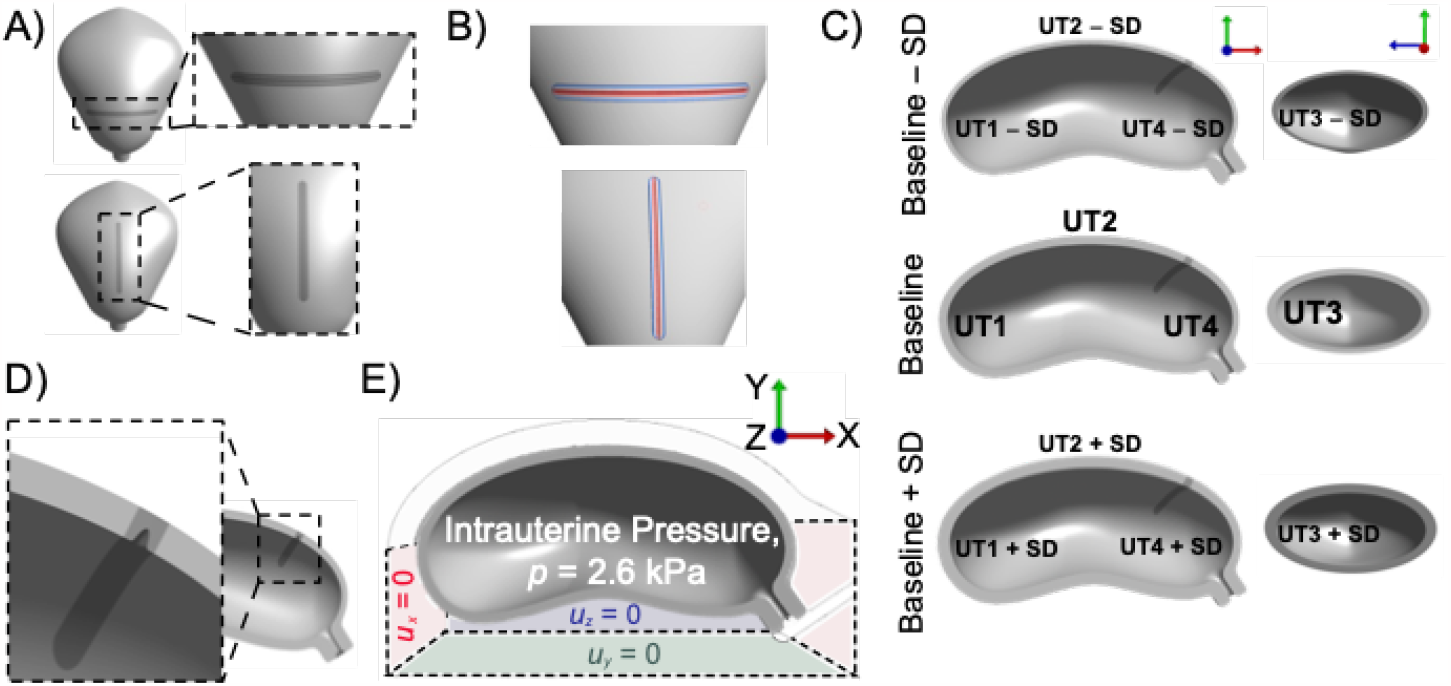
A) Representative baseline models of a uterus and cervix with sagittal and transverse C-section scars were generated from measurements acquired from MRI images of at-term pregnant patients. B) The scar geometry was split through the midline so that the stiffness (*E*) of the midline could be varied, and the remaining stiffness was linearly interpolated to the surrounding uterine stiffness. C) The uterine thickness was varied by increasing and decreasing the fundal (UT1), anterior (UT2), left/right (UT3) and lower uterine segment (UT4) thickness by a standard deviation (SD) of patient measurements. D) A defect was incorporated in the model with a low transverse scar. E) The displacement of the abdomen walls was fixed in the *x, y* and *z* directions and an intrauterine pressure was applied (*p* = 2.6 kPa).

### Mesh Generation

The abdomen, uterus, cervix, and scar geometries with and without defects were discretized into linear tetrahedral elements using Hypermesh 2022 (Altair, Michigan, U.S.A.). Results from our Baseline model with second-order tetrahedral elements were compared to results from our Baseline model with first-order elements and we found that the stress magnitudes were similar (the change in maximum von Mises stress was less than 1%). To reduce computation time, first-order elements were used. The element sizes were determined through a mesh convergence study to determine when the von Mises stress values (*σ*) of specified nodes in the uterus and scar geometries converged to a solution.

### Finite Element Simulation and Analysis

Meshed geometries were imported into FEBio Studio (v2.2) and modeled as a nearly incompressible (Poisson’s ratio *ν* = 0.48) neo-Hookean material according to the FEBio manual [21, 22]. The Young’s modulus (*E*) of the abdomen and the uterus were set to 100 kPa, while the stiffness of the scar tissue was varied. First, the scar tissue was treated as a homogeneous material and the Young’s modulus (*E*) of the scar was varied (50 kPa, 100 kPa, 200 kPa). Next, the Young’s modulus of the midline of the scar region was varied (50 kPa, 100 kPa, 200 kPa). The Young’s modulus of the remaining scar tissue was calculated by a linear interpolation between the midline value and the surrounding uterine tissue. Displacement of the abdomen walls in the *x, y* and *z* directions were fixed and a physiological intrauterine pressure (*p* = 2.6 kPa) was applied as shown in Fig. 1E [23]. The models were performed and analyzed using FEBio (v4.2) and FEBio Studio (v2.2). The von Mises stress was calculated as described in the FEBio Manual [24].

## 3 Results

To understand how the presence of a sagittal scar instead of a low transverse scar could increase the risk of uterine rupture, the von Mises stress on the scar region of both types of scars were compared. Heat maps of von Mises stress revealed that stress on the anterior side of the sagittal scar is greater than that on the anterior side of the transverse scar (Fig. 2A). The models in which the scar properties matched the surrounding tissue properties validated the boundary conditions and demonstrated the expected null-response. Additionally, varying the stiffness of the scar region from 50 kPa to 200 kPa showed that as the scar stiffness increases, so does the stress in the scar region, as expected (Fig. 2A). Plotting the von Mises stress in the scar region as a histogram normalized to element volume confirmed that the maximum stress was greater in the sagittal scar compared to the respective transverse scar with the same Young’s modulus (Fig. 2B). Next, the stiffness of the midline of the scar was varied to allow for a gradient change in stiffness to the scar midline. In this case, the results were similar to the homogeneous scars (Fig. 3).

**Fig. 2.**
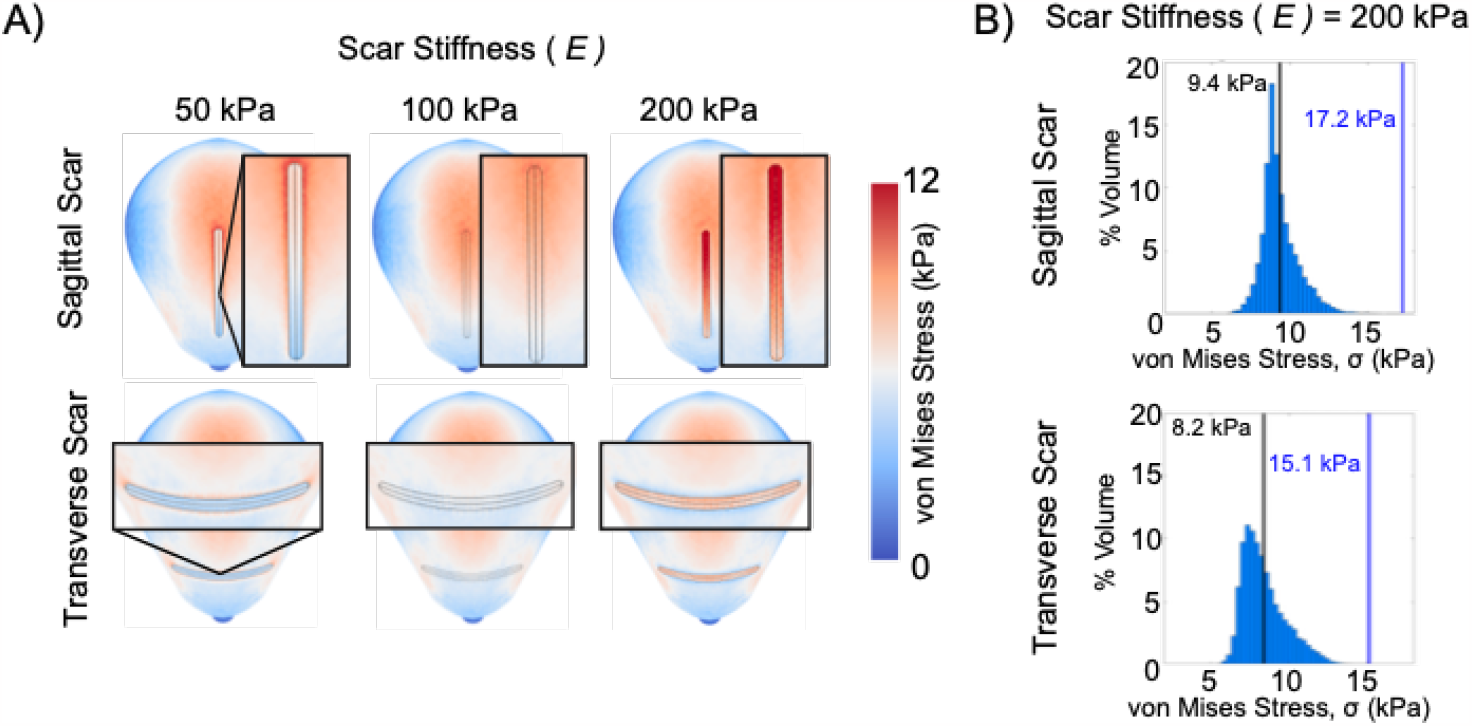
A) Heat maps of von Mises stress (*σ*) were generated and showed increased stress with sagittal scars compared to transverse scars with the same stiffness (*E*). B) The average (black line) and maximum (blue line) von Mises stress in the scar region of the sagittal scar was greater compared to the transverse scar with the same stiffness (*E* = 200 kPa).

**Fig. 3.**
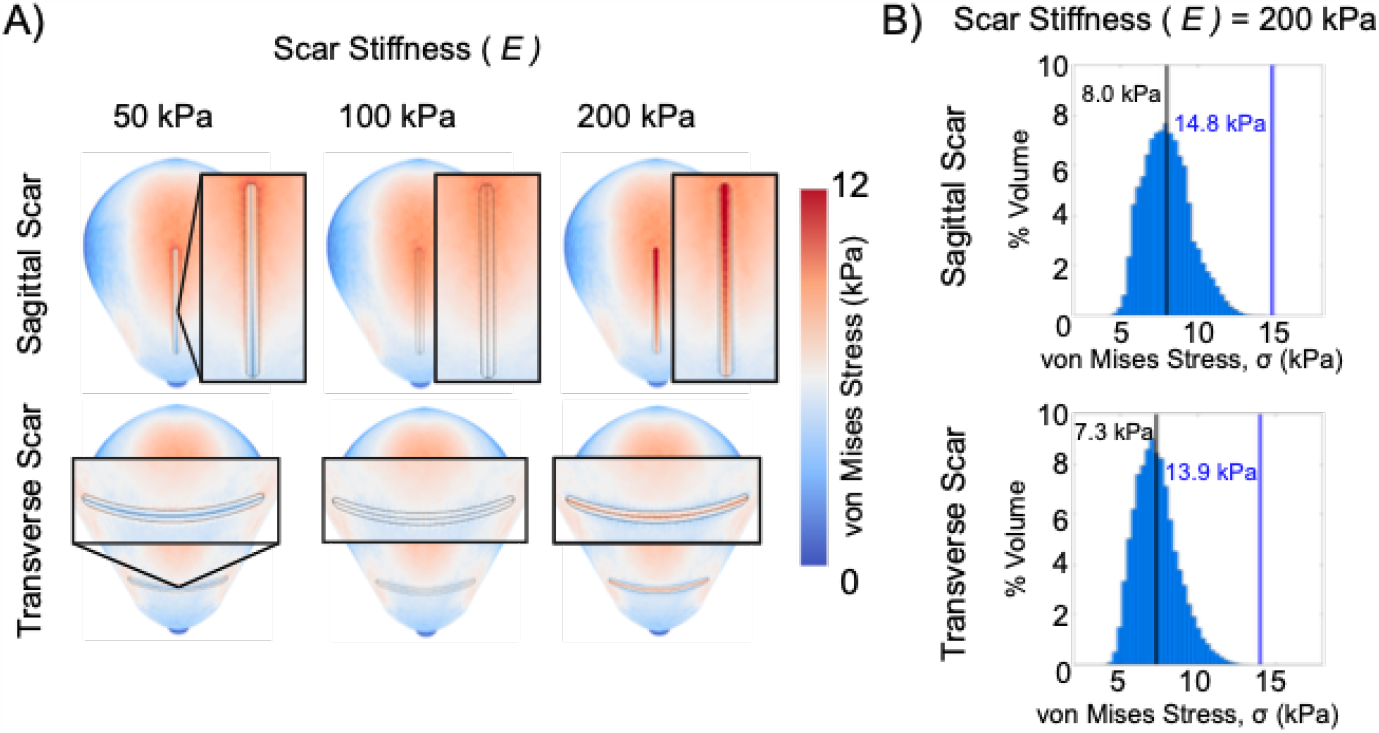
A) The stiffness (*E*) of the midline of the scar was varied, which allowed for a gradient of stiffness between the surrounding uterine tissue. Von Mises stress (*σ*) distributions show similar trends to the homogeneous scars. B) The average (black line) and maximum (blue line) von Mises stress in the scar region of the sagittal scar was greater compared to the transverse scar with the same stiffness (*E* = 200 kPa).

Since the thickness of the lower uterine segment has been shown to be predictive of uterine rupture, the thickness of the uterine tissue was also varied. When varying the thickness of the uterine wall, heat maps of the von Mises stress in the scar region showed that decreasing the wall thickness increased the stress in the scar region (Fig. 4). Plotting the calculated stress values of each element in the scar region as a histogram normalized to element volume confirmed that the mean and maximum stress does increase with decreasing thickness (Fig. 4).

**Fig. 4.**
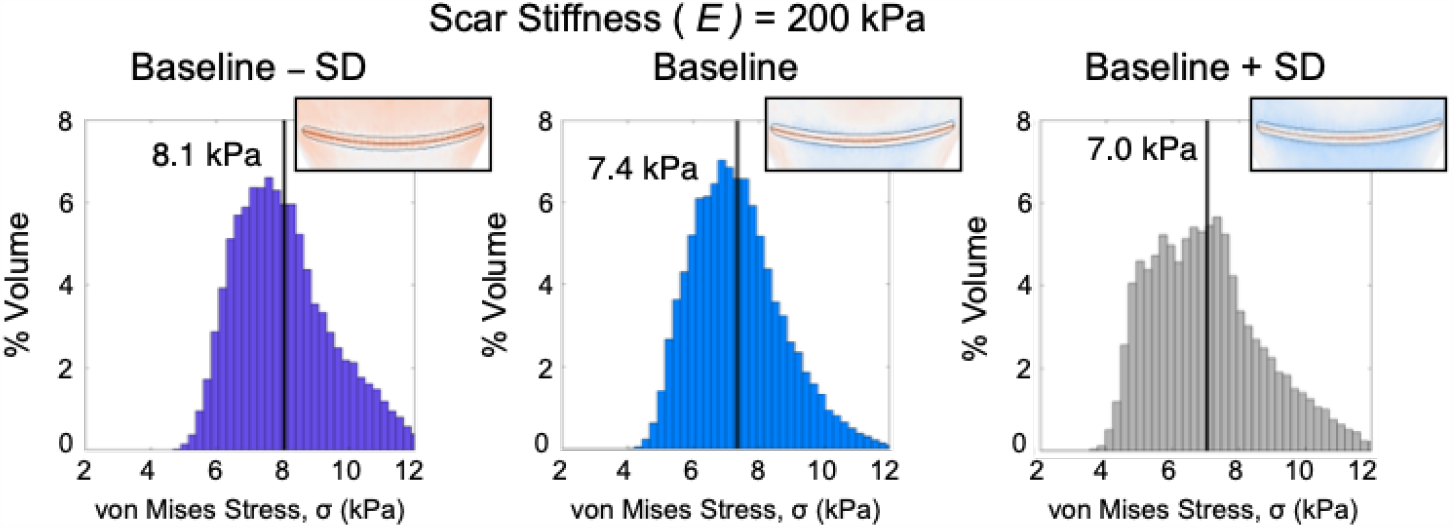
The uterine thickness was increased and decreased by the standard deviation (SD) of five patients’ uterine thickness measurements and the scar stiffness was held constant (*E* = 200 kPa). The mean von Mises stress (*σ*) in the scar region decreased with increasing uterine thickness.

Incorporating a defect in the low transverse scar caused stress concentrations in the posterior side of the defect region (Fig. 5A). Quantifying the stress along the midline of the scar from the outer edge to the center demonstrated that there is an increase in von Mises stress at the defect region compared to the model with no defect incorporated (Fig. 5B).

**Fig. 5.**
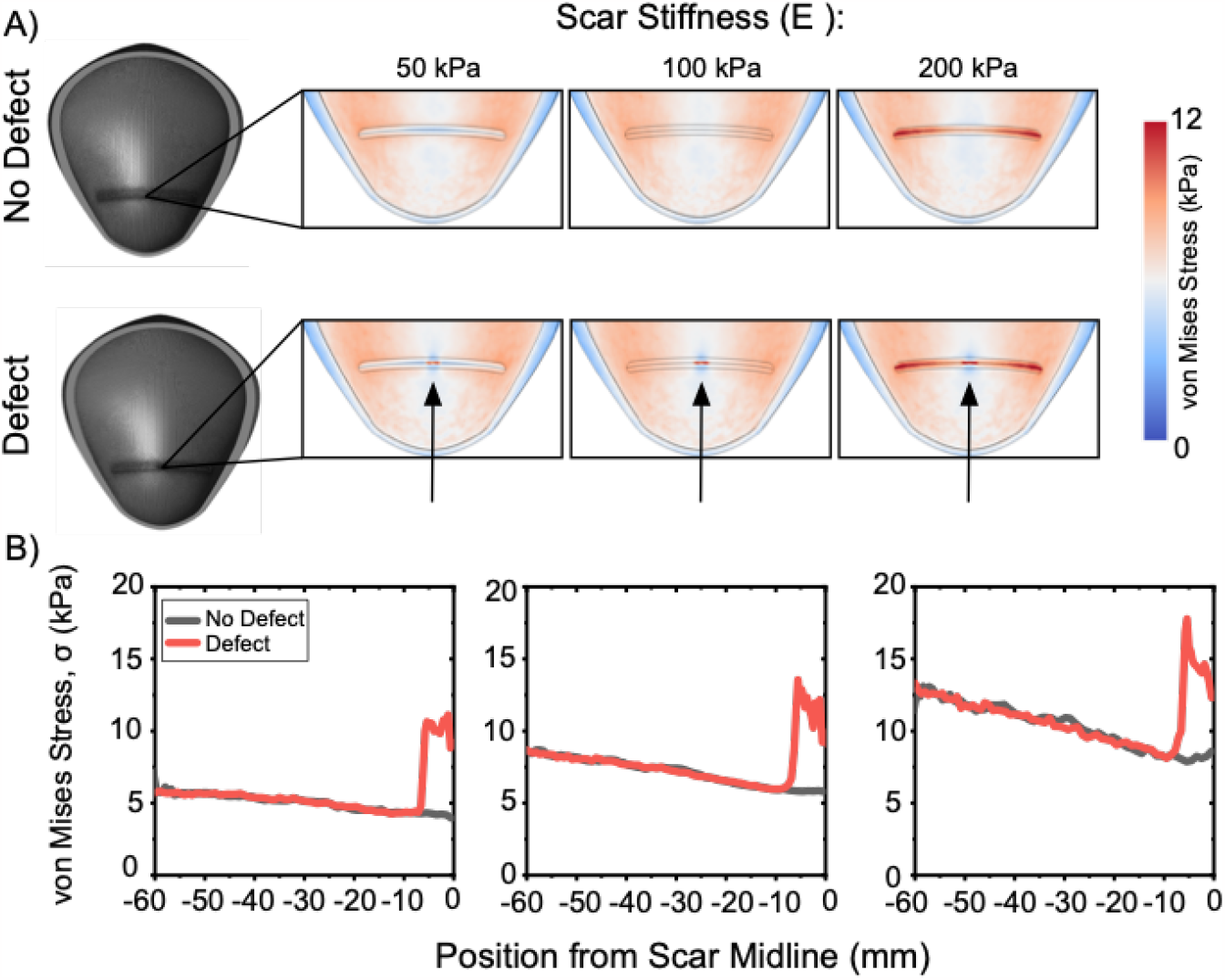
A) Compared to the models with no scar defect, the von Mises stress in the interior of the defect region increases. The stress on the defect also increases with increasing scar stiffness, as expected. B) Plotting the von Mises stress of elements along the edge of the scar from the outer edge (− 60 mm) to the center of the scar in the defect region (0 mm) demonstrates the increased stress in the defect region.

## 4 Discussion

In conclusion, scar positioning, decreasing uterine thickness, and the presence of a defect are all factors that increase the stress in the scar region, suggesting the tissue may be more likely to fail in this location. Taken together, the results of this proof-of-concept study are promising because the increased regions of stress due to scar positioning, thinning of the uterine walls and the presence of a defect are consistent with clinical observations of features that increase the risk of uterine rupture. While the results are promising, this work includes several limitations. The model assumes all tissues can be represented as a neo-Hookean material, but other materials, may be more representative of the uterine and scar tissue [25, 26]. For example, an anisotropic fibrous material would be more representative of the behavior of the uterus and C-section scar, especially since the fibrous nature of tissue generally increases with scar tissue because of the increased levels of collagen type I [25, 27]. Additionally, linear tetrahedral elements were used in this study, which are known to generate volumetric locking phenomena when used to simulate the response of incompressible materials, so future work will use higher order elements [28]. Also, a Young’s modulus range of 50 – 200 kPa was used, but more accurate measurements of material properties of uterine tissue could be incorporated into the model [29]. This model also did not account for spatial variation of material properties in the uterine tissue. While there was a spatial variation of material properties of the scar tissue, we assumed that the greatest change in the elastic modulus was at the midline of the scar tissue, which may not accurately represent the spatial variation of material properties in C-sections scars. Therefore, future work is needed to characterize the material properties and spatial variation of material properties of uterine tissue and C-section scars.

In addition to addressing limitations of this work, future research should also explore how models could predict fracture of uterine and C-section scar tissue to understand the mechanisms of uterine rupture. The fracture of highly deformable soft materials is an ongoing important field of research [30,31]. Developing predictive theories of soft material fracture is challenging due to the nonlinear and dissipative deformation involved [30]. Unlike typical engineering materials, small flaws can be present without leading to catastrophic failure and the stress-strain fields near the crack tip are nonlinear. Crack tips of soft materials can also exhibit crack-blunting and as a result highly deformable materials can resist cracks up to a few millimeters long, demonstrating excellent flaw tolerance. Additionally, as the model complexity is increased to include anisotropy of the tissue, von Mises yield criterion may not be relevant to failure of the material and other failure criterion may be more relevant [32].

Before modeling C-section scars and scar defects to predict risk of uterine rupture, it will be critical to follow the American Society of Mechanical Engineers (ASME) standards for verification and validation of solid mechanics models and models for medical devices [33]. Validating models and determining the reliability of models in reproductive health is extremely challenging since typical methods of validation using animal tissues are not an option because the animal physiology and anatomy of reproductive tissues differ from humans [34]. *Ex vivo* testing of tissues from hysterectomies may also be an option, however, results will still vary from *in vivo* situations. Therefore, future work will be an iterative process of refining the models, while comparing the outcomes to available clinical data.

Overall, this study demonstrates a framework to build patient specific geometries of C-section scars and scar defects to understand how these structures influence uterine deformation and stress under an applied load. In the future, further parametric models could identify critical parameters that influence uterine rupture risk and perhaps identify new assessments of risk that can be considered for patient specific surgical planning.

## Acknowledgements

The authors acknowledge the NIH T32 Postdoctoral Training Grant in Regenerative Medicine (T32 EB028092).

## Notes

### Competing Interest Statement

The authors have declared no competing interest.

### Summary of Updates

To correct typos found in the manuscript

